# Behavioral and physiological evidence that increasing group size ameliorates the impacts of social disturbance

**DOI:** 10.1101/818401

**Authors:** Hannah M. Anderson, Alexander G. Little, David N. Fisher, Brendan L. McEwen, Brett M. Culbert, Sigal Balshine, Jonathan N. Pruitt

## Abstract

Intra-group social stability is important for the long-term productivity and health of social organisms. We evaluated the effect of group size on group stability in the face of repeated social perturbations using a cooperatively breeding fish, *Neolamprologus pulcher*. In a laboratory study, we compared both the social and physiological responses of individuals from small versus large groups to the repeated removal and replacement of the most dominant group member (the breeder male). Individuals living in large groups were overall more resistant to instability but were seemingly slower to recover from perturbation. Members of small group were more vulnerable to instability but recovered faster. Breeder females in smaller groups also showed greater physiological preparedness for instability following social perturbations. In sum, we recover both behavioral and physiological evidence that living in larger groups helps to dampen the impacts of social instability in this system.

**Summary Statement:** Social stability is vital for group productivity and long-term persistence. Here, both behavioral and physiological evidence conveys that larger groups are less susceptible to social disturbance.

## Introduction

Living in groups has various costs and benefits. For instance, group living can increase foraging efficiency (Berger, 1978), decrease predation risk (Foster and Treherne, 1981), and increase collective reproductive output (Modlmeier et al., 2012). In contrast, living in groups can sometimes decrease average per capita reproductive output (Bilde et al., 2007), promote disease transmission (Kappeler et al., 2015), and increase competition for food (Symington, 1988). For group living to evolve, the weight of the combined benefits of grouping must exceed the costs, and any factor that maximizes benefits whilst minimizing the costs of living in groups should promote the evolution of group-living and help to optimize sociality once it has evolved.

One factor thought to help maximize the cost/benefit ratio of group living is social stability. For instance, increased familiarity among group members can allow for increased social niche specialization (Laskowski and Pruitt, 2014), reduced within-group competition (Laskowski and Pruitt, 2014), and increased group productivity (Modlmeier et al., 2012). Familiarity of groupmates can also enhance the effects of social buffering against environmental challenges (Hennessy et al., 2000; Livia Terranova et al., 1999) and decrease overall stress levels (Culbert et al., 2018; Kikusui et al., 2006; Nadler et al., 2016). Group stability also helps to reduce the costs of group living. For example, stable groups composed of familiar individuals experience less internal conflict, and so experience less stress from the threat of aggression or eviction (Pardon et al., 2004), reduced risk of injury, and waste fewer resources in competition (Marler, Walsberg, White, Moore, & Marler, 1995; Jordan et al 2010). Even in non-cooperative territorial species, familiarity among neighbors commonly begets reduced aggression via dear enemy effects (e.g., Getty, 1987; Siracusa et al., 2017).

Despite the common finding that group stability helps to maximize group success, all groups in nature must endure some level of instability. Immigration/emigration, birth/death, and alterations to dominance hierarchies, for example, result in alterations in group membership, and thus decrease within-group familiarity and stability. Many social species have therefore evolved mechanisms to help mitigate the negative impacts of such forces. For instance, some groups exhibit social rules that allow dominance hierarchies to swiftly reorganize following perturbation (Goldenberg et al., 2016). In other cases, reconciliatory communication mechanisms (e.g., specialized vocalization) aid in re-galvanizing damaged social bonds (Waal, 2000; Reddon et al 2019), and even particular individuals can help to dampen the negative impacts of group instability (Flack et al., 2005; Flack et al., 2006; McCowan et al., 2011). The traits that enable groups to dampen the acute impacts of social instability and to resume their former predictable states swiftly are important, because *i*) stabilizing traits are potentially important targets for selection and *ii*) forces that compromise these traits risk imperiling the integrity and function of the social system.

Here we examined how one group trait, group size, impacts the acute behavioral and physiological responses of group members to social disturbances and recoverability from them. We elected to focus on group size because it is known to mediate many costs and benefits associated with group living (Avilés and Tufino, 1998), and because natural groups vary considerably in their size, with profound impacts on social selection (Brown et al., 2016). We predicted that living in large groups would diminish the acute impacts of social perturbations and increase group recoverability by distributing the negative impacts of social disturbance (e.g., acts of aggression) across more individuals. Larger groups may also recover more swiftly via enhanced affiliative behavior following social perturbations. We term this the ***distributed perturbation hypothesis*** here. Alternatively, living in larger groups might increase the negative impacts of social perturbations (e.g., via increased aggression) or prevent groups from resuming quiescent behavioral states following disturbance. For instance, aggressive acts might initiate positive feedback fostering additional aggressive interactions in high-density environments and thus prevent groups from resuming their former stable states. We term this the ***aggressive feedback hypothesis***.

The impacts of social disturbances are likely to be evidenced physiologically as well. We therefore evaluated whether group size alters the degree to which group members are metabolically poised for intense bouts of acute or sustained physical activity following social perturbation. A higher capacity for intense activity might be necessary in preparation for, or as a training effect of, increased aggression. Many studies have identified links between various social behaviors and metabolic rates (see Huntingford, Tamilselvan, & Jenjan, 2012 for review). However, reliance on oxygen consumption as a proxy for energy metabolism neglects the anaerobic processes that fuel burst-type movements typically associated with dominance behaviors (Plaut, 2001). Thus, a more refined focus on the biochemical pathways that underlie metabolic phenotypes should help elucidate links between physiology and behaviour.

Enzymes are catalytic proteins that regulate biochemical reaction rates (Boyer and Krebs, 1986). Their expression is often plastic and can change in response to environmental stressors over a period of days to weeks (Beaman et al., 2016). Enzymes that catalyze regulatory steps of greater biochemical pathways can thus be plastically adjusted to meet an organism’s peak metabolic demands in contrasting environments. Thus, *in vitro* measures of regulatory enzyme activities can represent upper thresholds for their respective pathways, and reflect the maximum capacity for these pathways to fuel peak activity *in vivo* (e.g.Vigelsø, Andersen, & Dela, 2014). Indeed, a number of studies have shown that activities of specific metabolic enzymes correlate strongly with intense social behaviors in a range of animal systems (Gilmour et al., 2017; Guderley, 2009; Guderley and Couture, 2005; Kasumovic and Seebacher, 2013; Le François et al., 2005; Regan et al., 2015). In this study we focused on a key regulatory glycolytic enzyme (lactate dehydrogenase; LDH) and a key regulatory oxidative enzyme (citrate synthase; CS) that have been shown to reflect capacities for quick burst movements and more sustained aerobic activities, respectively (e.g. Alp, Newsholme, & Zammit, 1976; Childress & Somero, 1979; Johnston & Moon, 1981). We hypothesized that LDH and CS activities would scale with the most intense bouts of dominant actions displayed by an individual, and that these activities would be highest in individuals from destabilized groups.

To address these questions, we use the cooperative breeding cichlid *Neolamprologus pulcher*, endemic to Lake Tanganyika in the African Rift Valley. In the wild groups are usually comprised of one dominant male-female breeding pair and 1-20 smaller, subordinate, non-breeding helpers (Balshine et al., 2001; Heg et al., 2019). Groups cooperate to care for the young of the dominant pair, maintain the group’s territory, and defend the territory from both competitors and predators (Taborsky & Limberger, 1981; Wong & Balshine, 2011a). These fish also have a clear linear size-based dominance hierarchy, with increasing body size associated with increasing rank (Balshine-Earn et al., 1998). Natural groups regularly experience turnover in group members as helpers join or leave a group, or when group members perish (Stiver et al 2006; 2007; Heg et al., 2019; Wong & Balshine, 2011b). Thus, this system provides a convenient evolutionary context to evaluate the impacts of group size on behavioral and metabolic responses to social instability and recoverability.

## Methods

### Ethics

All experimental protocols were approved by the Animal Research Ethics Board of McMaster University (Animal Utilization Protocol No. 18-04-16), and were in compliance with the guidelines set by the Canadian Council on Animal Care (CCAC) regarding the use of animals in research.

### Behavioral Methods

Focal fish were haphazardly selected from a lab population containing descendants of wild-caught *N. pulcher* captured in 2014. Large and small groups were formed with a dominant pair (the largest male and female in each social group), and either four (“small groups”, n=12) or eight (“large groups”, n=14) subordinate helper fish. To reduce aggression and mortality, dominant pairs were taken from pre-existing breeding pairs. All helpers were unfamiliar to the dominant pair and had not previously cohabitated with them. Following group formation, the social groups were allowed to habituate and stabilize for five weeks.

Each social group was maintained in separate, 189 L aquariums containing two terracotta pot halves and two small PVC tubes (that served as both shelter and breeding substrate), two 10 cm x 10 cm mirrors, two sponge aeration filters, a heater, and 3 cm deep coral sand as substrate. The mirrors served as a target of aggression to reduce morbidity from within-group conflict. A water temperature of 27° C and 13:11 light to dark hour photoperiod was maintained throughout the study. Each dominant male and female received an identifying dorsal fin clip, which has a minimal effect on behavior (Stiver et al., 2004). Fish were fed six days a week ad libitum with Nutrafin® basix large cichlid flakes.

Small and large social groups were randomly allocated to either control (large, n=6; small, n=6) or treatment (large, n=8, small, n=6) conditions. The dominant male (standard length, SL: average=7.57 cm, SEM=0.92) and dominant female (SL: average=6.66 cm, SEM=0.86) were measured at the start of the experiment. To avoid confusion with later measures, these fish will subsequently be referred to as the *breeding male* and *breeding female*. The standard lengths of all helpers were estimated by an experienced observer (SB) (SL: average=2.52 cm, SEM=0.07). In the treatment condition, the social perturbation consisted of the removal of the breeding male from one social group and replacing him with a new, unfamiliar breeding male that previously dwelled in another social group of identical size, tank set up, and group composition. Therefore, breeding male fish in the treatment groups were swapped between tanks. We ensured that the breeding males were always larger than the females, as observed in the wild (Balshine et al., 2001; Desjardins et al., 2008; Wong et al., 2012)In the control condition, the breeding male fish were removed from their tanks, handled for the same duration as the treatment males, but then returned to their home tank. This social disturbance procedure occurred twice (trial 1 and trial 2), with the manipulations conducted one week apart. All tanks were perturbed on the same day. Perturbations were conducted twice to permit group members time to deploy an enzymatic response to reliably stable vs. perturbed social conditions.

Behavioral observations were recorded using Canon VIXIA HF r-series cameras immediately before the manipulation, immediately following the manipulation, and then four, and twenty-four hours following the manipulation. In addition, two baselines were recorded twenty-four and forty-eight hours prior to the first manipulation. Focal observation recordings were all fifteen minutes long. The first five minutes of each recording were discarded to account for potential disturbance on remaining group members from capturing and returning the dominant male fish and human presence. All videos were scored by the same observer (HA), who was blind to treatment condition and time step. Behaviors were coded using McMaster University’s Animal Behavior Ecology Laboratory (ABEL) *N. pulcher* ethogram (Sopinka et al., 2009) and Behavioral Observation Research Interactive Software (BORIS) (Friard and Gamba, 2016). Behaviors were subdivided into the categories, “aggression” (chase, bite, ram, puffed throat, mouth-fighting, pseudo-mouth-fighting and head shake)), “submission”, (submissive posture, submissive display, flee/chased and bitten)) and “affiliation” (soft touch, following, and parallel swim).

We calculated a Dominance Index (DI) for each breeding male, breeding female, and for each group of helpers divided per capita, for each recording session. The DI is a well-established method for calculating dominance rank and = (sum of aggressive acts given + sum of submissive acts received) - (aggressive acts received + submissive acts given). We calculated an affiliation index for each breeding male, breeding female, and for each group of helpers divided per capita, for each recording session, where affiliation rank and = (sum of social acts given + sum of social acts received). We also recorded the most dominant time step for breeding females in each tank, herein referred to as the *maximum dominance index observed*. Specifically, the maximum dominance index observed represents the DI of the time period with the highest levels of aggressive and submissive behaviors. This term therefore reflects what are presumably the most stressful and metabolically demanding moments we observed (Grantner and Taborsky, 1998).

The breeding female of each group was captured and rapidly (≤3 minutes) euthanized via overdose of benzocaine within forty-eight hours of the final perturbation. Females were measured and their skeletal muscle just posterior to the dorsal fin, heart, and liver were harvested and massed for further analyses.

### Enzyme Assays

In short, tissues were homogenized in 1:10 (m/v) homogenization buffer (0.1% Triton, 50 mM Hepes, 1mM EDTA, pH 7.4; CAT: 100 mM K phosphate buffer, 100 mM KCl, 1 mM EDTA, pH 7.4) on ice. Skeletal muscle homogenates were further diluted to 1:400 for the LDH activity assay, whereas liver homogenates were diluted to 1:20 for both LDH and CS activity assays. Skeletal muscle homogenates were not further diluted for CS activity assays. All assays were run at 27°C in 96-well format on a Spectramax Plus 384 microplate reader (Molecular Devices, Sunnyvale, CA). We used a wavelength of 340 nm to measure the disappearance of NADH (for LDH activity), and a wavelength of 412 nm to measure the production of 2-nitro-5-thiobenzoic acid (TNB; as a proxy of CS activity). Extinction coefficients of 6.22 mM-1 cm-1 and 13.6 mM-1 cm-1 were used for LDH and CS, respectively.

### Analyses and Statistical Methods

Dominance and affiliation indices were analyzed using a mixed linear model (GLMM) fit by REML using the free and open software JAMOVI (Version 0.9; GAMLj module; https://www.jamovi.org). We fitted tank number as a random effect, and focal individual class (i.e. female, male, helpers), treatment type (i.e. control vs. treatment), group size, trial number (i.e. trial 1 or trial 2), and timepoint (i.e. immediately before the manipulation, immediately after, four hours after and twenty-four hours after the manipulation) as fixed effects. We started with a saturated model and pruned non-significant terms (starting with high-order interactions) until we arrived at a model where all fixed effects were significant. *Post-hoc* analyses consisted of Bonferroni-corrected pairwise comparisons.

To analyze the relationship between body traits (mass, relative heart mass, liver mass) and the maximum dominance index observed during the experiment on metabolic capacity (glycolytic and aerobic) across females, we used general linear models (GLM) fit by OLS. For the maximum dominance index observed, we fitted treatment type and group size (factors) and body mass, relative heart mass, and liver mass (continuous covariates) as fixed effects. For metabolic capacity, LDH activity in either the muscle or the liver, or CS activity in either the muscle or the liver represented the dependent variable. Treatment type, group size (factors), maximum dominance index observed, body mass, and other enzyme activity levels (continuous covariates) were fitted as fixed effects. We used the maximum dominance index observed as a fixed effect because LDH and CS measures convey individuals’ capacities for peak activity. Thus, in addition to generalized locomotor activity these effects also likely determine maximum capacities for social activities (e.g. aggression, flight, and dominance), rather than baseline averages. We again started with a saturated model and pruned non-significant terms (starting with high-order interactions) until we arrived at a model where all fixed effects were significant. As a *post-hoc* approach to test whether the effects of maximum dominance on enzyme activities were a potential effect of activity levels, we fitted respective models using mean activity measures as a covariate in place of maximum dominance. For all statistical tests, we used a significance threshold of α = 0.05.

## Results

### Behavioral responses

We detected a significant four-way interaction between class, treatment type, group size, and time point on individuals’ dominance scores (Table 1; Fig. 1A-D). In control tanks housing small groups, male dominance was consistently more than five-fold greater that of females, although this trend was significant only immediately after the control perturbation (Fig. 1A; S1 for pairwise comparisons). In control tanks housing large groups, there were no significant differences in dominance between the males, females, and helpers; although the helpers consistently had a five-fold lower dominance scores than both the males and females (Fig. 1B; S1). These results suggest that male aggression is more pronounced in small control groups and that females display more submissive acts in response.

**Table 1.** Statistical parameters for final (minimal) GLMM for dominance and affiliation indices.

| **Dominance** | **Fixed Factor** | **F** | **Num df** | **Den df** | **p** |
| --- | --- | --- | --- | --- | --- |
|  | Treatment | 5.439 | 1 | 22 | 0.029 |
|  | Time point | 1.137 | 3 | 554 | 0.334 |
|  | Group Size | 8.138 | 1 | 22 | 0.009 |
|  | Class | 146.405 | 2 | 554 | < .001 |
|  | Treatment * Time point | 0.44 | 3 | 554 | 0.724 |
|  | Treatment * Group Size | 4.782 | 1 | 22 | 0.04 |
|  | Time point * Group Size | 0.514 | 3 | 554 | 0.673 |
|  | Treatment * Class | 31.266 | 2 | 554 | < .001 |
|  | Time point * Class | 3.944 | 6 | 554 | < .001 |
|  | Group Size * Class | 4.463 | 2 | 554 | 0.012 |
|  | Treatment * Time point * Group Size | 0.527 | 3 | 554 | 0.664 |
|  | Treatment * Time point * Class | 3.809 | 6 | 554 | < .001 |
|  | Treatment * Group Size * Class | 2.242 | 2 | 554 | 0.107 |
|  | Time point * Group Size * Class | 5.069 | 6 | 554 | < .001 |
|  | Treatment * Hours * Group Size * Subject | 3.685 | 6 | 554 | 0.001 |
| **Affiliation** |  |  |  |  |  |
|  | Group Size | 0.00442 | 1 | 22 | 0.948 |
|  | Treatment | 3.12959 | 1 | 22 | 0.091 |
|  | Trial # | 11.43825 | 1 | 580 | < .001 |
|  | Class | 8.02106 | 2 | 580 | < .001 |
|  | Time point | 1.98344 | 3 | 580 | 0.115 |
|  | Group Size * Treatment | 1.25136 | 1 | 22 | 0.275 |
|  | Treatment * Time point | 1.75654 | 3 | 580 | 0.154 |
|  | Trial # * Time point | 5.4469 | 3 | 580 | 0.001 |
|  | Group Size * Time point | 1.1949 | 3 | 580 | 0.311 |
|  | Group Size * Treatment * Time point | 2.74729 | 3 | 580 | 0.042 |

Numerator degrees of freedom, Num df; Denominator degrees of freedom, Den df; Shading, p < 0.05.

**Figure 1.**
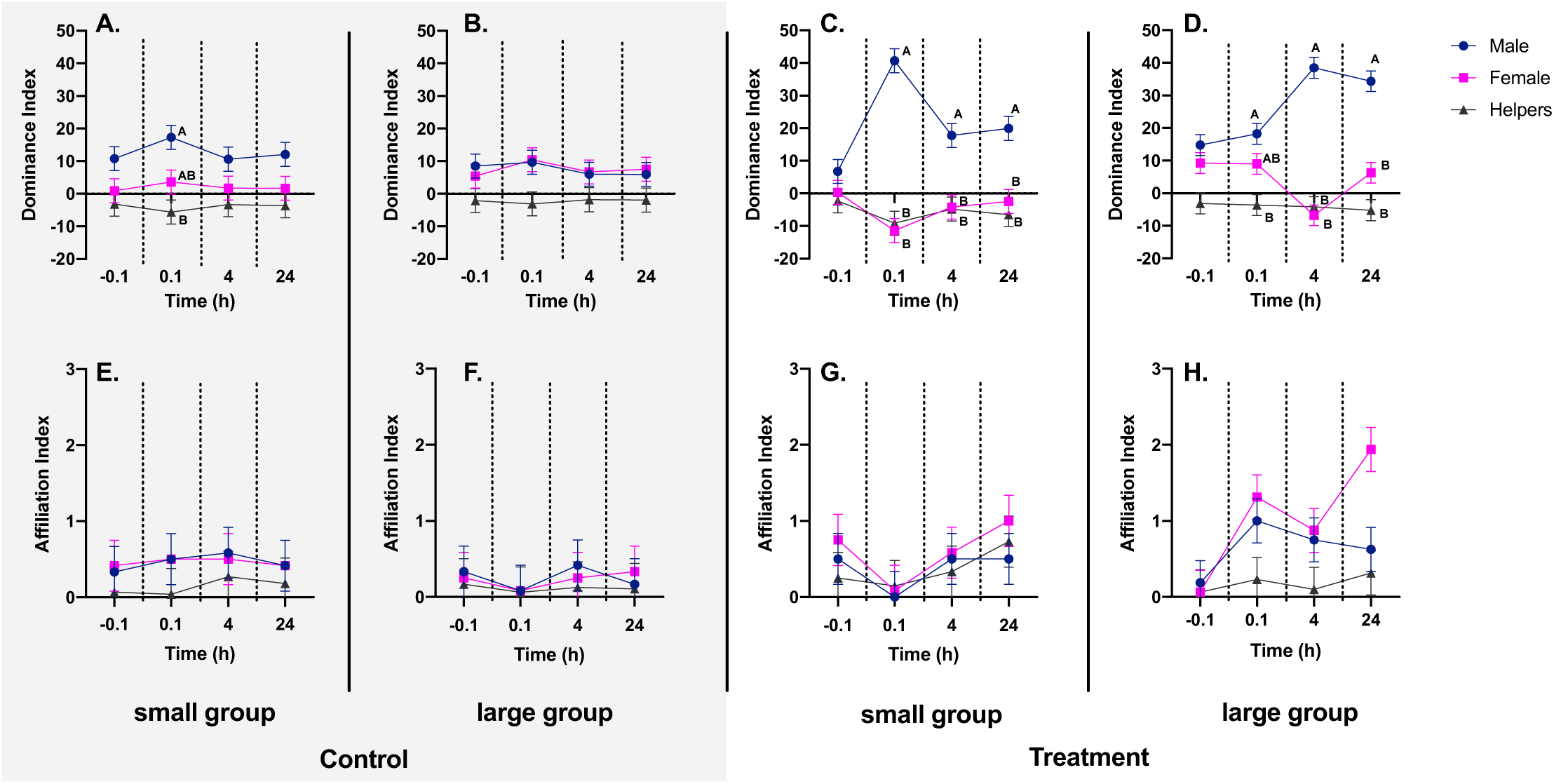
Time courses for mean dominance and affiliation indices. Mean dominance indices in males (blue), females (pink), and helpers (grey) from small and large control (A and B, respectively) and treatment (C and D, respectively) groups. Mean affiliation indices from small and large control (E and F, respectively) and treatment (G and H, respectively) groups. Different letters represent differences between males, females and helpers within each respective timepoint, as determined by *post hoc* comparisons.

In treatment tanks housing small groups, we found that the dominance indices of the females were significantly lower than that of the males at all time points, especially immediately following the perturbation (Fig. 1C; S1). However, in treatment tanks housing large groups there was a delayed spike in male dominance relative to females, where no significant difference in dominance between males and females was apparent until four-hours after the perturbation (Fig. 1D; S1). As expected, helper dominance remained significantly lower than male dominance across all time points in both group sizes. There was no significant effect of trial number (i.e. perturbation 1 vs. perturbation 2) in any of the analyses.

There was a significant interaction term between group size, treatment, and time point on social affiliation scores. We further detected a significant interaction term between trial number and time point, and a main effect of individuals’ class (female, male, helper) on social affiliation scores (Table 1; Fig. 1E-H). While there was no effect of group size on affiliation scores in the control groups, affiliation conspicuously increased following perturbation in the large treatment groups relative to the small treatment groups. Groups gradually increased affiliative behaviors following the introduction of a new male, but somewhat decreased affiliative behavior following the introduction of a second new male (Table 1; see appendix for pairwise comparisons). Finally, females had the highest affiliation index followed by males, and then by helpers in the treatment groups (Fig. 1E-H; Table 1; S2 for pairwise comparisons).

We found an interaction between body mass and group size on the maximum dominance index observed (Table 2; Fig. 2a). Here, maximum scores for dominance increased with female body size in small groups and decreased with body size in large groups. There were no significant effects of relative heart or liver size on maximum scores for dominance, and no main effect of treatment type (i.e. control vs. treatment).

**Table 2.**
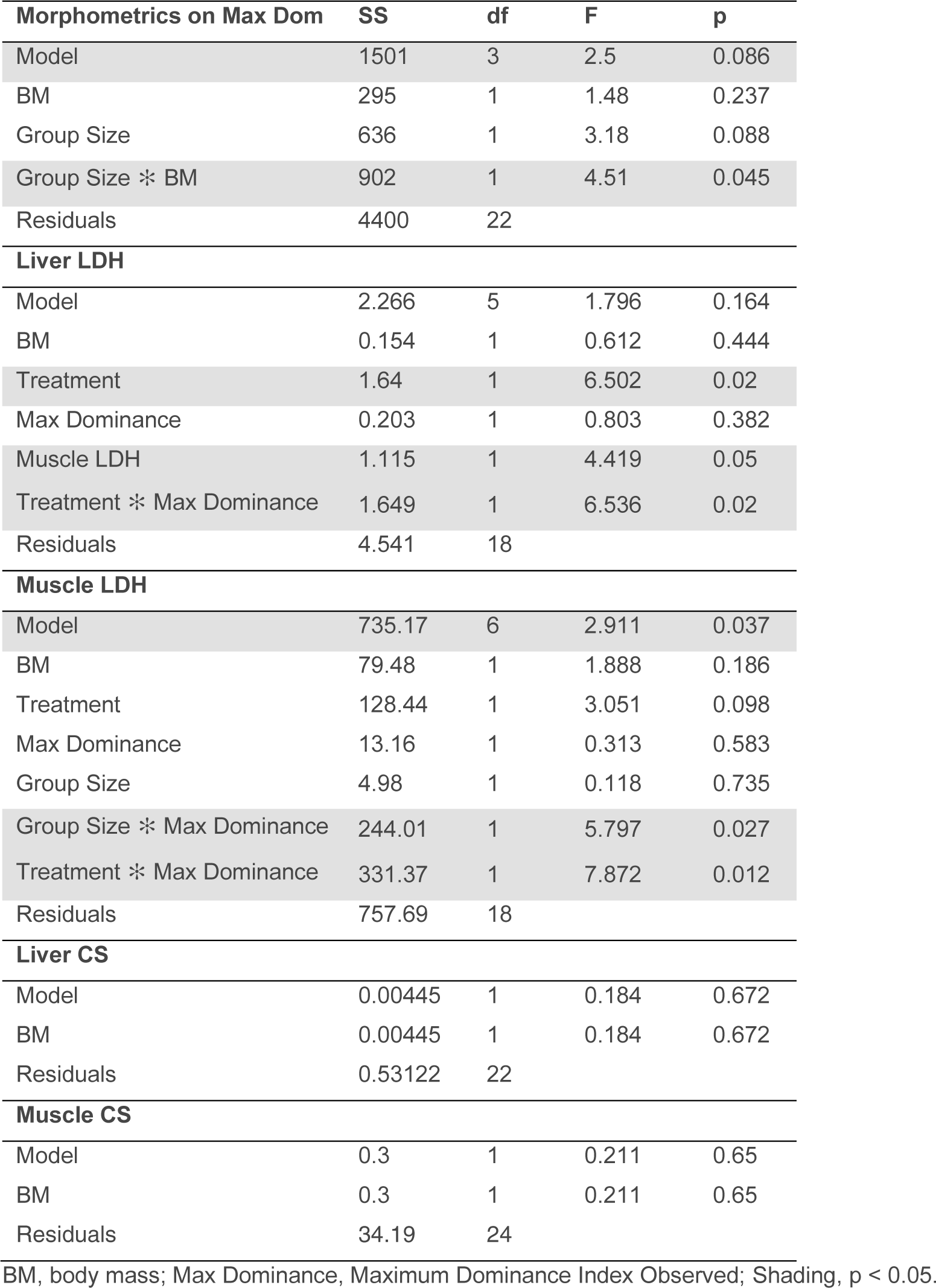
Statistical parameters for final (minimal) GLM for female-level effects.

**Figure 2.**
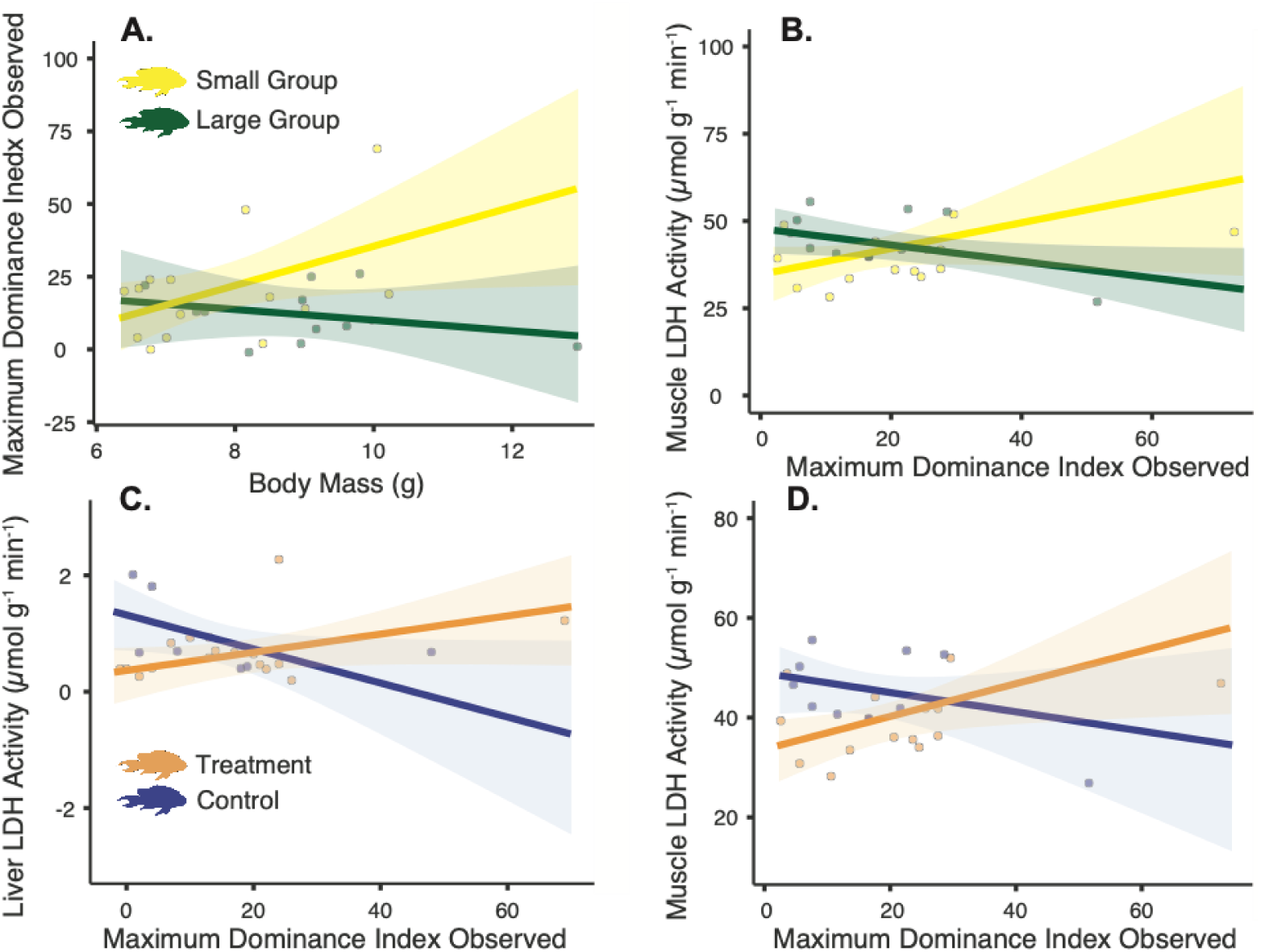
Relationships between the maximum dominance we observed (Maximum Dominance Index) and body mass (A), Muscle LDH activity (B), Liver LDH activity (C), and Muscle LDH activity (D). Small and large groups (A,B) are represented by yellow and green, respectively, whereas treatment and control groups (C,D) are represented by orange and blue, respectively. Enclosed circles represent observed scores. Note, the directionality and patterns of the relationship remain when we remove the two most extreme data points.

### Enzyme responses

There was a significant interaction between female dominance and group size on muscle LDH activity (Table 2; Fig. 2B). Muscle LDH activity scaled positively with dominance in small groups and it scaled negatively with dominance in large groups, suggesting that breeding females were more poised for intense bursts of activity in smaller groups. We also found that both liver and skeletal muscle LDH activities scaled negatively with female dominance in control groups, and positively with female dominance in treatment groups (Table 2; Fig. 2C, D). These results convey that our social perturbation treatment was successful in causing the breeding females to be enzymatically prepared for sudden bursts of activity. We found no significant effects of dominance on liver or muscle CS activity (Table 2), which is associated with endurance activities and aerobic metabolism. Instead, the anaerobic component of metabolism as captured by LDH activity was more responsive to our social perturbations. *Post-hoc*, we found no significant effects of mean level of female activity on liver or skeletal muscle LDH activities (appendix 4), suggesting that maximum dominance affects glycolytic capacity independently from greater levels of general locomotor activity.

## Discussion

Group stability tends to increase the benefits and decrease the costs of social living (Berger, 1978; Laskowski and Pruitt, 2014; Modlmeier et al., 2012), and groups often exhibit mechanisms to return to a stable state following disturbance (Goldenberg et al., 2016; McCowan et al., 2011; Waal, 2000). We sought to determine the effects of group size on the group’s ability to return to social homeostasis in the face of a repeated social stressor. Specifically, we hypothesized a large group would either reduce overall aggression, through the *distributed perturbation hypothesis*, or increase and sustain overall aggression, through the *aggressive feedback hypothesis*. Here we found more support for the *distributed perturbation hypothesis*, though additional moderating forces are also likely at play.

Small groups showed more disparate dominance indices between the most dominant fish (breeding males) and the subordinate fish (breeding females and helpers). This is most obvious when comparing the control groups (Fig. 1A,B). Previous studies have found a positive relationship between group size and long-term group survival (Heg et al., 2019), with large groups benefitting from higher quality territories and more opportunities to feed (Balshine et al., 2001). These latest results further imply that small groups may be inherently more polarized (and less stable) than large groups, even when social conditions remain relatively steady. In other words, large groups likely benefit from both material and non-material social advantages. The timing of dominance index spikes varied with group size in our treatment groups: in small groups, changes to and inequality of dominance indices appeared immediately following the perturbation (Fig. 1C), while in large groups change in the indices lagged following perturbation (Fig. 1D). Small groups also appear to slide back towards baseline states faster, as observed in the apparent reduction in breeding male dominance twenty-four hours following the perturbations, while the dominance of large group males remain elevated. Together, these results suggest that large groups are more resistant to social state change and/or that state change in large groups is slower than in small groups. This could be because new males delay asserting their dominance in larger groups until they have had time to evaluate their new social setting and potential competitors. Regardless of the mechanism, this conveys that larger groups might offer their constituents buffering effects against ephemeral social perturbations in a manner small groups do not.

Additional circumstantial evidence from affiliation indices and body mass hint that smaller groups are more stressful social environments following perturbation. One can observe an increase in the affiliative behaviors of males and especially females following social perturbations in large groups (Fig. 1H). This conveys that the new breeding pair begins establishing a social bond in these groups. If this happens in small groups too, then it is certainly less conspicuous (Fig. 1G). We further note that large females exhibit higher dominance in small groups, irrespective of control vs. treatment, whereas no relationship between body size and dominance was observed in large social groups. This group-size dependent relationship conveys that more volatile acts of dominance transpire in small groups occupied by large females, whereas the dominance indices of females in large groups are near uniformly low (Fig. 2A). This lack of variation in large groups provides further evidence that large social groups are less volatile and more stable social environments than small groups. In *N. pulcher* the strength of social buffering is largely managed by aggression rates (Culbert et al., 2019), so the decreased aggression found in these large groups might facilitate recovery from social perturbation. Elevated LDH activities in muscle and liver suggest enhanced glycolytic preparedness and capacity for the powerful burst movements that characterize aggressive acts (Le François et al., 2005). In Arctic charr (*Salvelinus alpinus*), for instance, fast-twitch muscle fibers of dominant individuals possess LDH activities more than 15% greater than their subordinate counterparts (Le François et al., 2005). Our work, however, shows that group size directionally mediates the relationship between dominance and glycolytic capacity. LDH activity was highest in the most dominant animals but only in small groups, which also have the most disparate dominance indices between males and females (Fig. 1A, C). In large groups however, more dominant females were characterized by lower LDH activity levels. These trends suggest that the more dominant females in small groups must be better primed to perform (or potentially avoid) aggressive actions, while the more dominant females in large groups are not. Whether these phenotypic differences reflect a regulated response to social stress, a positive feedback effect of training, or a combination of the two, remains to be examined. However, the lack of relationship between these enzyme measures and greater female activity levels suggests these trends are not simply a feedback effect of exercise training. Together, our findings suggest that breeding females in small groups experience greater instability following disturbance and are metabolically prepared for more instability.

The divergent relationship between dominance and LDH activity provides evidence that our social perturbations were successful in instigating an enzymatic response in females. Muscle and liver LDH activities increased with female dominance in treatment groups, which were characterized by the largest gaps in dominance between males and females. This further suggests that female dominance increases metabolic preparedness for aggression in these groups relative to controls. By contrast, in the control condition, LDH activity levels decreased with female dominance, suggesting greater dominance is associated with reduced glycolytic capacity and potentially greater stability in these groups. Because the control perturbation was characterized by a familiar male, we suggest that preestablished social relationships dampen the aggressive actions that foster glycolytic capacity.

Overall, we found more support for the *distributed perturbation hypothesis* from both behavioral and physiological indicators. Physiologically, breeding females elevated their glycolytic capacity in small groups and when faced with strong social perturbations (treatment). Behaviorally, small groups also showed a larger difference in dominance indices across group members, while in large groups’ dominance indices were slower to polarize following a perturbation and were associated with a surge of affiliative behaviors as well, both observations circumstantially supporting the *distributed perturbation hypothesis*. On the other hand, the gap in dominance indices shrunk faster following the perturbation in small groups compared to large, potentially supporting the *aggressive feedback hypothesis*. It therefore appears that different group sizes create different responses to the forces of instability: small groups experience larger instability following a social perturbation, recover more rapidly but are physiologically primed for more instability, whereas large groups are more resistant to the instability of perturbation but appear to recover more slowly. In aggregate, these results convey that the demographic traits of social groups can play a large role in shaping group susceptibility to and recoverability from social disturbance and that larger groups could exhibit greater levels of social stability and social inertia.

## Acknowledgements

Funding for these studies herein was generously provided by the Natural Sciences and Engineering Research Council of Canada Discovery Grants to JNP and SB and the Canada 150 Research Program to JNP.

## Data Availability

Behavioral data will be uploaded upon completion as supplementary.

---

**File S1**. Post-hoc pairwise comparisons for dominance index GLMM.

**File S2**. Post-hoc pairwise comparisons for affiliation index GLMM.

**Appendix 1.** Statistical parameters for the saturated GLMM for dominance indices.

|  | F | Num df | Den df | p |
| --- | --- | --- | --- | --- |
| Treatment | 5.4276 | 1 | 528 | 0.020 |
| Perturbation | 9.96e-4 | 1 | 528 | 0.975 |
| Time Point | 1.1063 | 3 | 528 | 0.346 |
| Group Size | 8.1205 | 1 | 528 | 0.005 |
| Subject | 142.4906 | 2 | 528 | < .001 |
| Treatment * Perturbation | 0.3960 | 1 | 528 | 0.529 |
| Treatment * Time Point | 0.4287 | 3 | 528 | 0.733 |
| Perturbation * Time Point | 0.0933 | 3 | 528 | 0.964 |
| Treatment * Group Size | 4.7720 | 1 | 528 | 0.029 |
| Perturbation * Group Size | 3.23e-4 | 1 | 528 | 0.986 |
| Time Point * Group Size | 0.5007 | 3 | 528 | 0.682 |
| Treatment * Subject | 30.4299 | 2 | 528 | < .001 |
| Perturbation * Subject | 2.5110 | 2 | 528 | 0.082 |
| Time Point * Subject | 3.8386 | 6 | 528 | < .001 |
| Group Size * Subject | 4.3436 | 2 | 528 | 0.013 |
| Treatment * Perturbation * Time Point | 0.0426 | 3 | 528 | 0.988 |
| Treatment * Perturbation * Group Size | 0.3010 | 1 | 528 | 0.583 |
| Treatment * Time Point * Group Size | 0.5125 | 3 | 528 | 0.674 |
| Perturbation * Time Point * Group Size | 0.1396 | 3 | 528 | 0.936 |
| Treatment * Perturbation * Subject | 2.6552 | 2 | 528 | 0.071 |
| Treatment * Time Point * Subject | 3.7067 | 6 | 528 | 0.001 |
| Perturbation * Time Point * Subject | 0.8327 | 6 | 528 | 0.545 |
| Treatment * Group Size * Subject | 2.1819 | 2 | 528 | 0.114 |
| Perturbation * Group Size * Subject | 0.6660 | 2 | 528 | 0.514 |
| Time Point * Group Size * Subject | 4.9333 | 6 | 528 | < .001 |
| Treatment * Perturbation * Time Point * Group Size | 0.0948 | 3 | 528 | 0.963 |
| Treatment * Perturbation * Time Point * Subject | 0.5351 | 6 | 528 | 0.782 |

|  | **F** | **Num df** | **Den df** | **p** |
| --- | --- | --- | --- | --- |
| Treatment * Perturbation * Group Size * Subject | 1.9841 | 2 | 528 | 0.139 |
| Treatment * Time Point * Group Size * Subject | 3.5868 | 6 | 528 | 0.002 |
| Perturbation * Time Point * Group Size * Subject | 0.5734 | 6 | 528 | 0.752 |
| Treatment * Perturbation * Time Point * Group Size * Subject | 0.8689 | 6 | 528 | 0.517 |

Numerator degrees of freedom, Num df; Denominator degrees of freedom, Den df.

**Appendix 2.** Statistical parameters for the saturated GLMM for Affiliation indices.

Numerator degrees of freedom, Num df; Denominator degrees of freedom, Den df. **Appendix 2.** Statistical parameters for the saturated GLMM for Affiliation indices.
|  | **F** | **Num df** | **Den df** | **p** |
| --- | --- | --- | --- | --- |
| Group Size | 0.00442 | 1 | 22.0 | 0.948 |
| Treatment | 3.12960 | 1 | 22.0 | 0.091 |
| Trial # | 10.75636 | 1 | 506.0 | 0.001 |
| Subject | 6.41299 | 2 | 506.0 | 0.002 |
| Hours | 1.93144 | 3 | 506.0 | 0.124 |
| Group Size * Treatment | 1.25136 | 1 | 22.0 | 0.275 |
| Group Size * Trial # | 1.15353 | 1 | 506.0 | 0.283 |
| Treatment * Trial # | 1.17067 | 1 | 506.0 | 0.280 |
| Group Size * Subject | 0.49489 | 2 | 506.0 | 0.610 |
| Treatment * Subject | 1.57884 | 2 | 506.0 | 0.207 |
| Trial # * Subject | 0.83349 | 2 | 506.0 | 0.435 |
| Group Size * Hours | 1.16357 | 3 | 506.0 | 0.323 |
| Treatment * Hours | 1.71051 | 3 | 506.0 | 0.164 |
| Trial # * Hours | 4.70137 | 3 | 506.0 | 0.003 |
| Subject * Hours | 0.62838 | 6 | 506.0 | 0.708 |
| Group Size * Treatment * Trial # | 0.16617 | 1 | 506.0 | 0.684 |
| Group Size * Treatment * Subject | 2.01673 | 2 | 506.0 | 0.134 |
| Group Size * Trial # * Subject | 0.71484 | 2 | 506.0 | 0.490 |
| Treatment * Trial # * Subject | 0.98941 | 2 | 506.0 | 0.373 |
| Group Size * Treatment * Hours | 2.67528 | 3 | 506.0 | 0.047 |
| Group Size * Trial # * Hours | 1.04450 | 3 | 506.0 | 0.372 |
| Treatment * Trial # * Hours | 1.98722 | 3 | 506.0 | 0.115 |
| Group Size * Subject * Hours | 0.57562 | 6 | 506.0 | 0.750 |
| Treatment * Subject * Hours | 0.36084 | 6 | 506.0 | 0.904 |
| Trial # * Subject * Hours | 0.95883 | 6 | 506.0 | 0.453 |
| Group Size * Treatment * Trial # * Subject | 0.59126 | 2 | 506.0 | 0.554 |
| Group Size * Treatment * Trial # * Hours | 0.26749 | 3 | 506.0 | 0.849 |
| Group Size * Treatment * Subject * Hours | 0.49915 | 6 | 506.0 | 0.809 |
| Group Size * Trial # * Subject * Hours | 0.28290 | 6 | 506.0 | 0.945 |
| Treatment * Trial # * Subject * Hours | 0.52073 | 6 | 506.0 | 0.793 |
| Group Size * Treatment * Trial # * Subject * Hours | 0.42951 | 6 | 506.0 | 0.859 |

Numerator degrees of freedom, Num df; Denominator degrees of freedom, Den df.

**Appendix 3.**
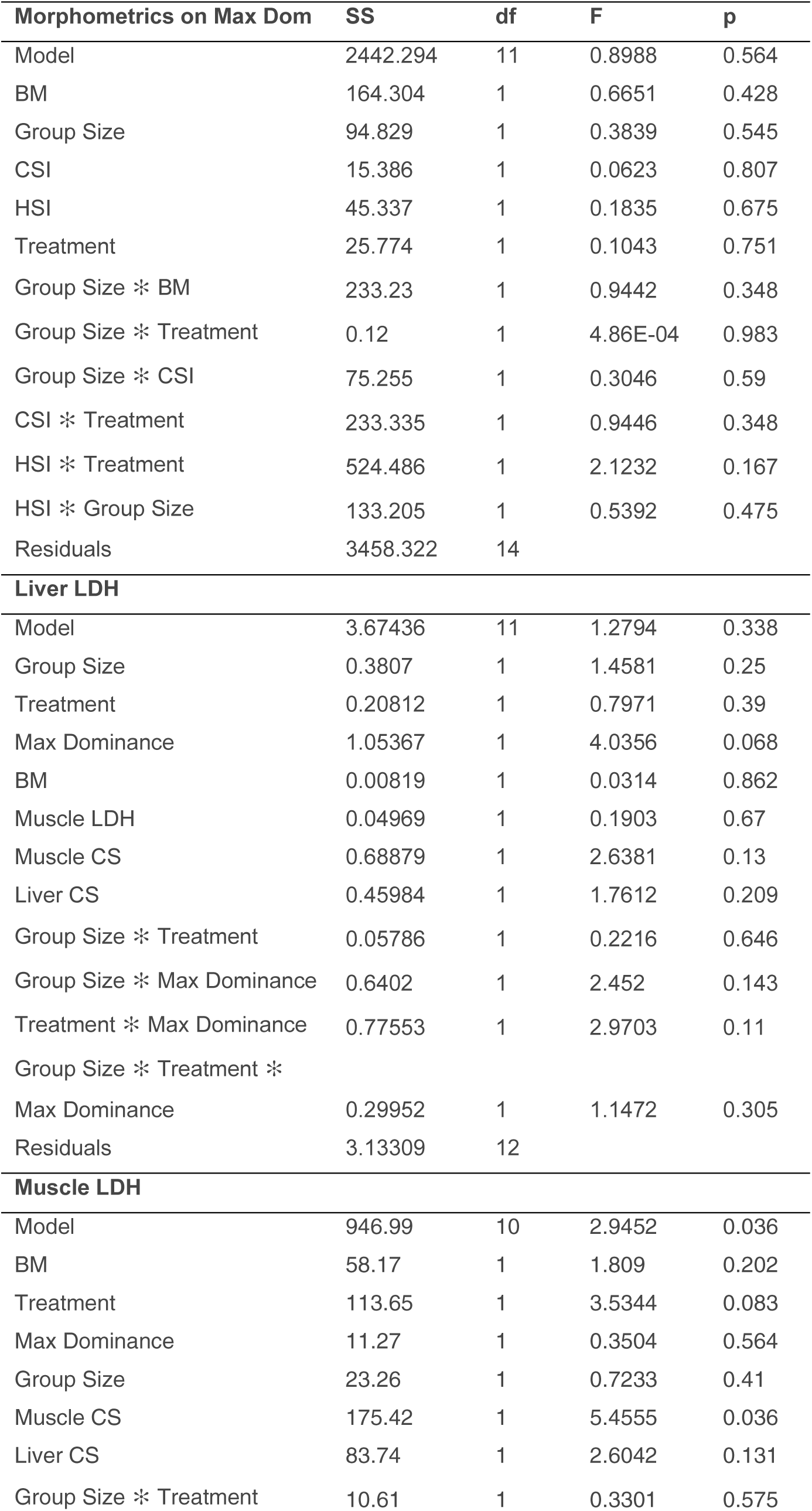
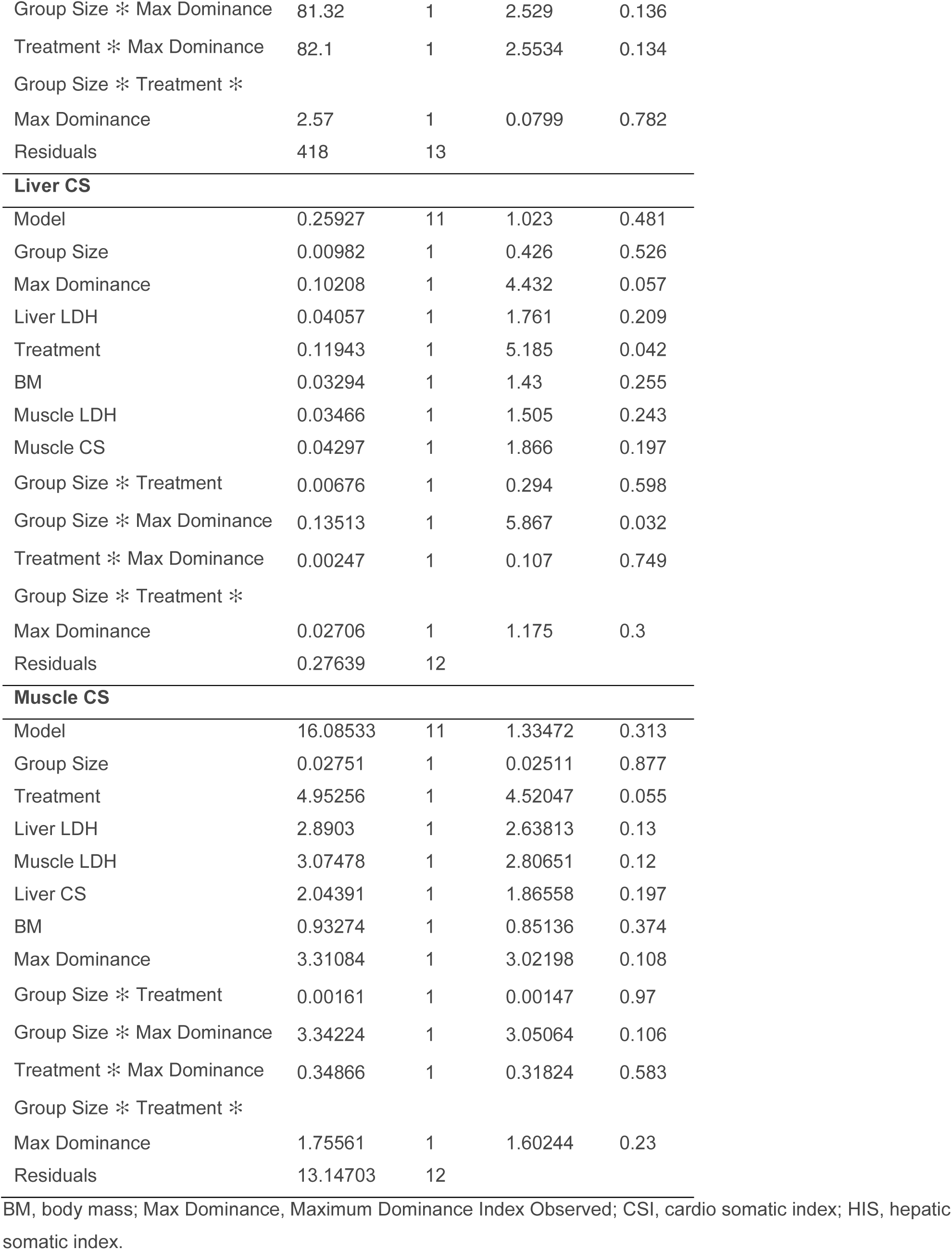
Statistical parameters for saturated GLMs for female-level effects.

**Appendix 4.** Statistical parameters for final (minimal) GLM for female-level effects.

| **Liver LDH** |  |  |  |  |
| --- | --- | --- | --- | --- |
| Model | 0.841 | 5 | 0.507 | 0.767 |
| BM | 0.196 | 1 | 0.593 | 0.451 |
| Treatment | 0.118 | 1 | 0.356 | 0.558 |
| Muscle LDH | 0.263 | 1 | 0.794 | 0.385 |
| Mean Activity | 0.131 | 1 | 0.394 | 0.538 |
| Treatment * Mean Activity | 0.173 | 1 | 0.522 | 0.479 |
| Residuals | 5.967 | 18 |  |  |
| **Muscle LDH** |  |  |  |  |
| Model | 286.556 | 6 | 0.71265 | 0.644 |
| BM | 37.637 | 1 | 0.5616 | 0.463 |
| Treatment | 105.763 | 1 | 1.57816 | 0.225 |
| Group Size | 7.757 | 1 | 0.11575 | 0.738 |
| Mean Activity | 46.458 | 1 | 0.69323 | 0.416 |
| Treatment * Mean Activity | 0.127 | 1 | 0.00189 | 0.966 |
| Group Size * Mean Activity | 27.164 | 1 | 0.40534 | 0.532 |
| Residuals | 1206.302 | 18 |  |  |

BM, body mass; Max Dominance, Maximum Dominance Index Observed; Shading, $p < 0.05$ .

